## Supplementary material for "Literate programming for iterative design-build-test-learn cycles in bioengineering"

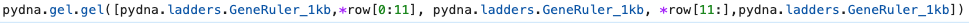

A)
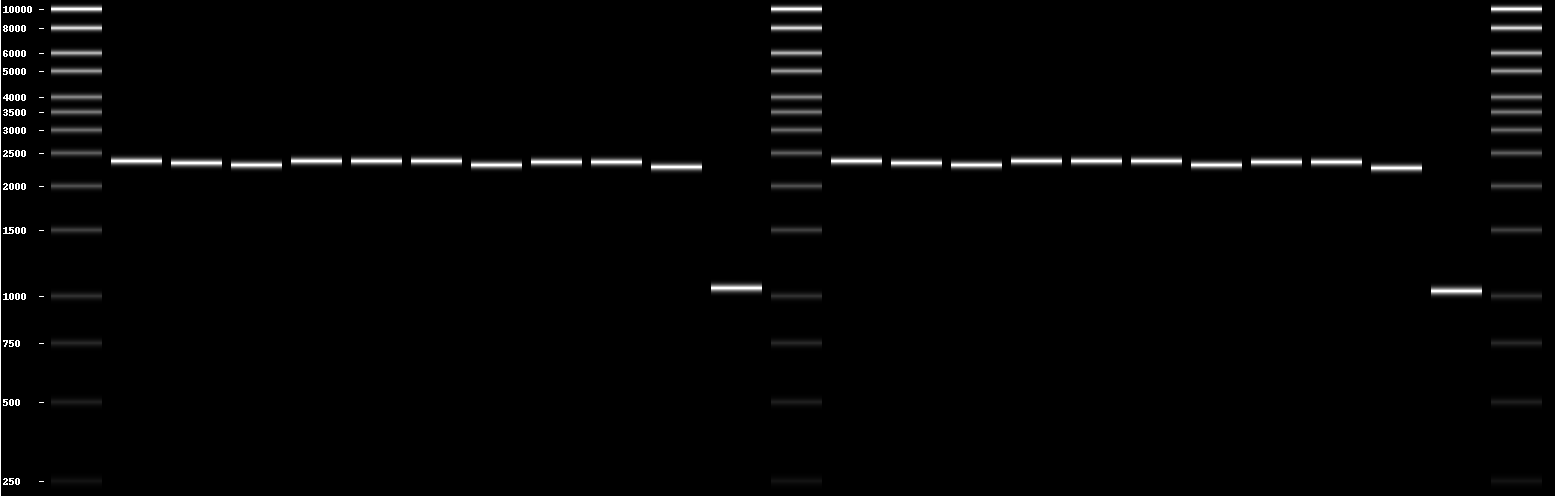

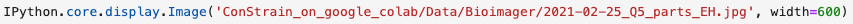

B)
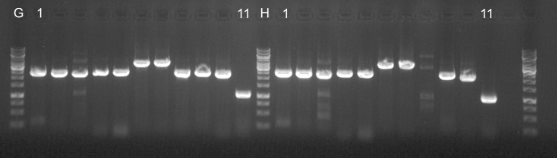

**Suppl. Fig S1.** Using literate programming to A) Simulate a gel with one line of code and B) Run the gel with the amplicons.

A)
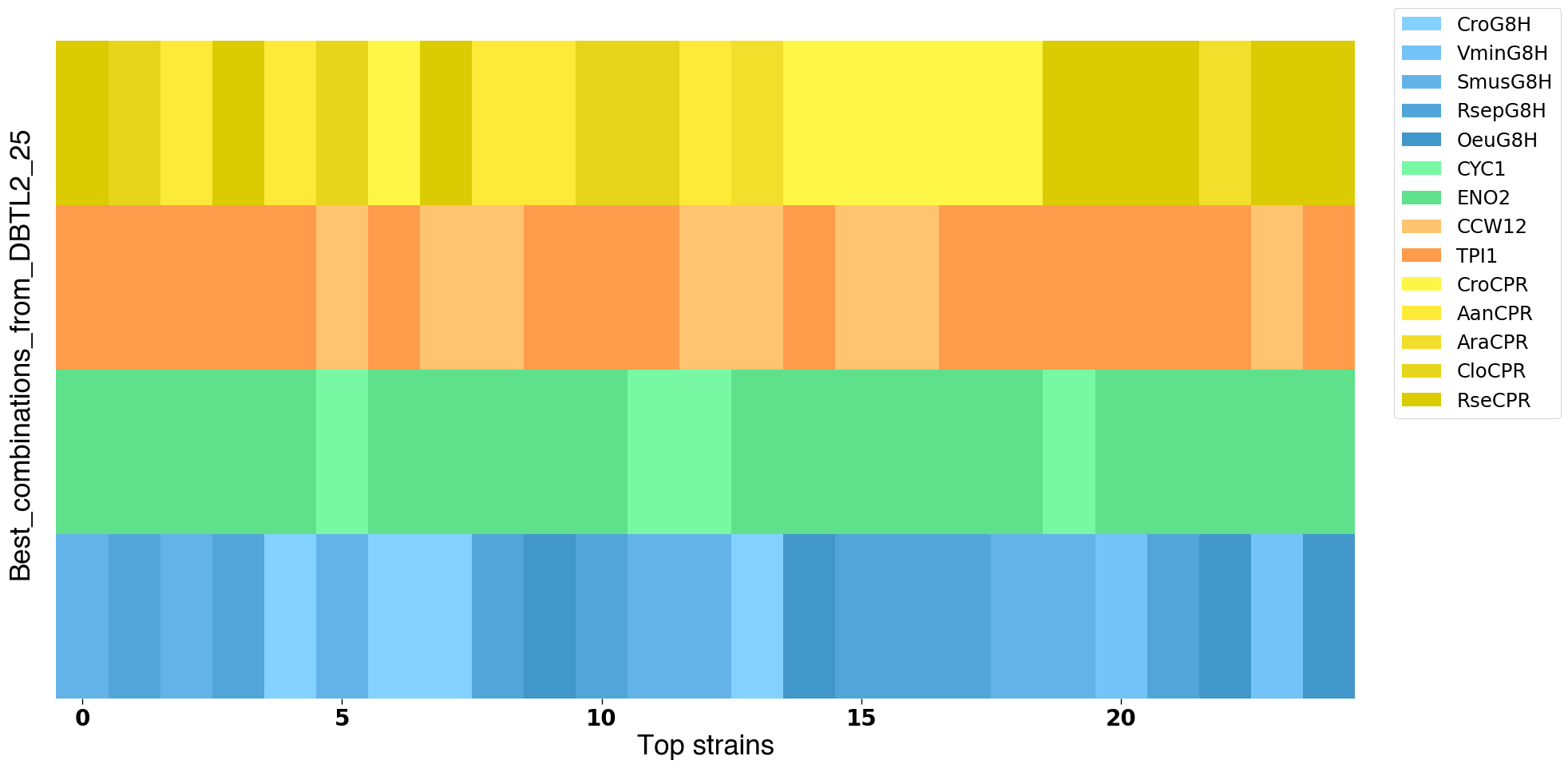

B)
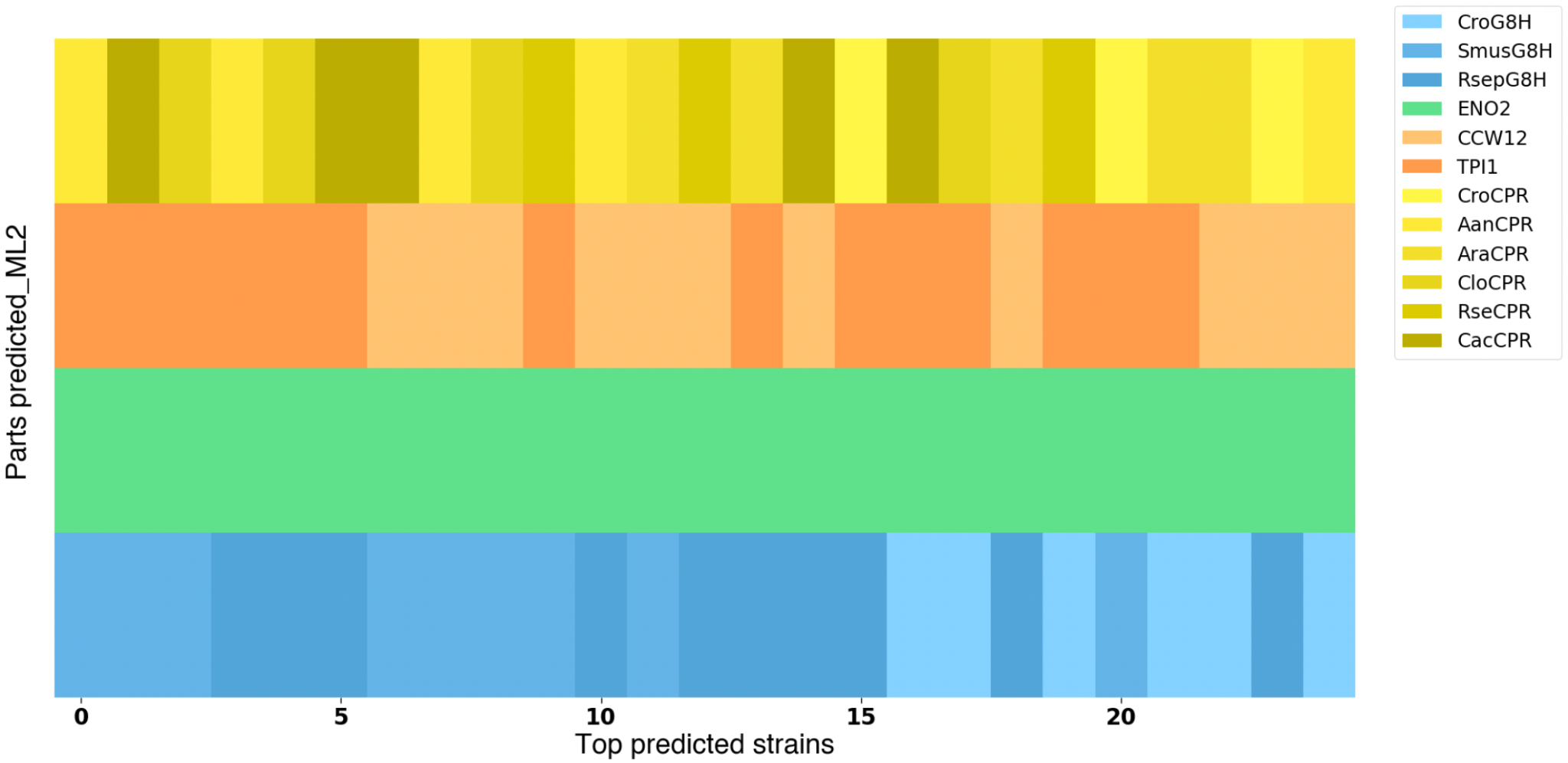

**Suppl. Fig S2.** Schematic representations of A) top 25 designed strains from DBTL2 and B) top 25 predicted strains from the updated machine-learning model after DBTL2.

C)
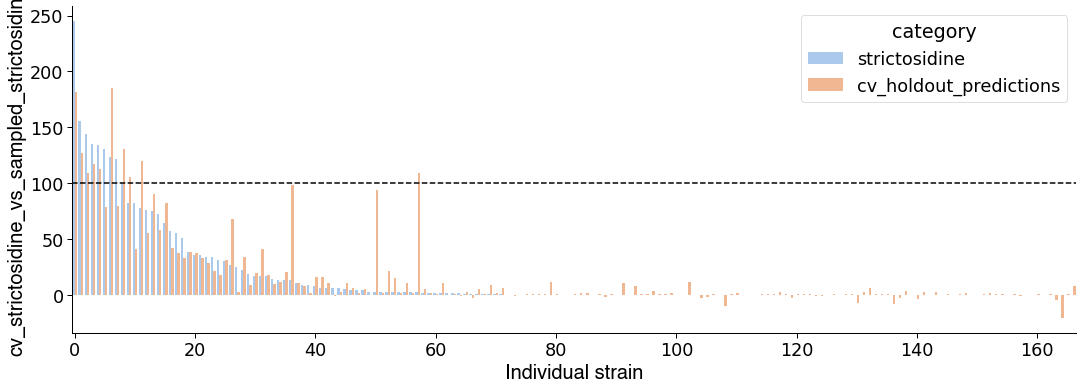

D)
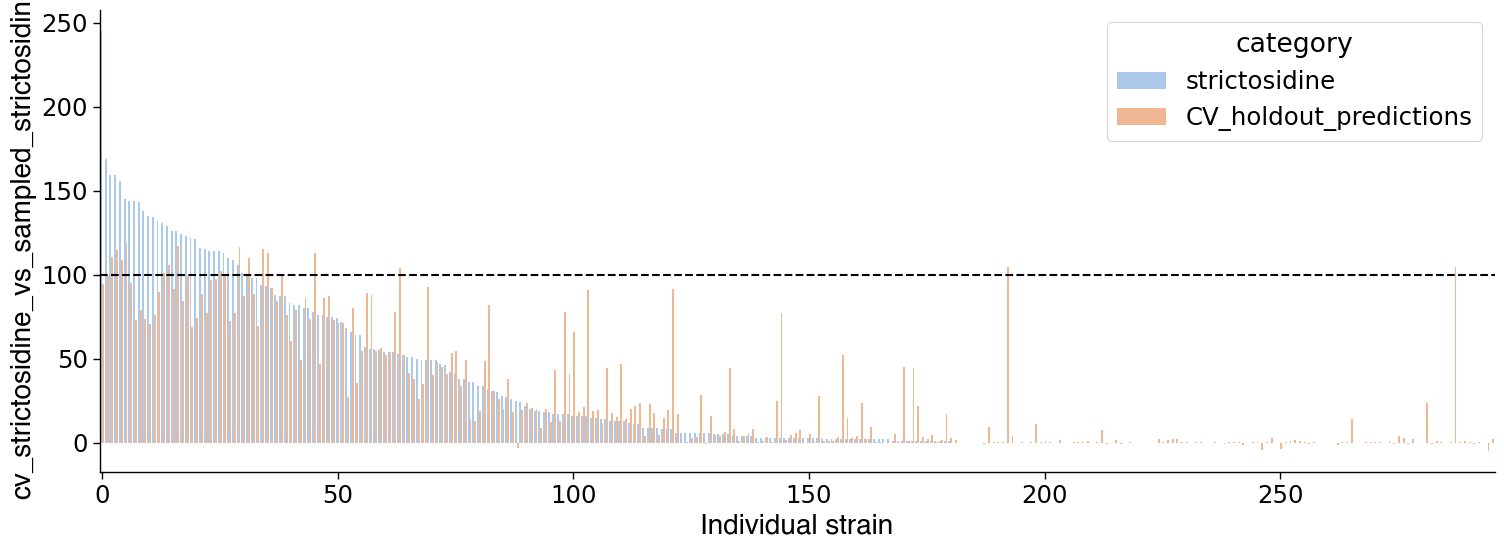

**Suppl. Fig S3.** A) Showing the observed strictosidine production values vs. the the cross-validated values from the model in the (A) first DBTL cycle and B) second DBTL cycle with all accepted strains.

**Table 1. Shows the dependencies for teemi divided into three categories: Minimal, Test, and Extra dependencies.**

| **Minimal Dependencies** | **Test Dependencies** | **Extra dependencies** |
| --- | --- | --- |
| pydna>=4.0.7 | pytest==7.1.2 | intermine==1.12.0 |
| pandas>=1.3.0 | pylint==2.13.9 | dnachisel==3.2.8 |
| benchlingapi>=2.1.12 | black==22.3.0 |  |
| numpy>=1.21.0 | pytest-cov==3.0.0 |  |
| biopython>=1.79 |  |  |
| python-dotenv>=0.20.0 |  |  |
| openpyxl>=3.0.9 |  |  |
| wheel>=0.37.1 |  |  |
| nbsphinx==0.8.9 |  |  |

**Table 2. Sorted strain performance of all fully genotyped strains in the first DBTL round.**

| **plate-well** | **names** | **Amt_uM_Loganin** | **Amt_uM_Secologanin** | **Amt_uM_Tryptamine** | **Amt_uM_Strictosidine** | **normalized_strictosidine(%)** |
| --- | --- | --- | --- | --- | --- | --- |
| yp50_D03 | SmusG8H-pENO2-pTPI1-AraCPR | 26,23584712 | 0 | 2021,748028 | 85,21014946 | 245,0342747 |
| yp50_G03 | RsepG8H-pENO2-pTPI1-RseCPR | 8,632278805 | 0 | 1476,111166 | 54,36248186 | 156,3272849 |
| yp50_E05 | SmusG8H-pCYC1-pTPI1-CacCPR | 8,009680311 | 0 | 1731,472157 | 50,19422696 | 144,340857 |
| yp51_A02 | OeuG8H-pENO2-pTPI1-CroCPR | 5,086582304 | 0 | 1798,561547 | 46,44331945 | 135,6934975 |
| yp50_A09 | VminG8H-pENO2-pTPI1-AraCPR | 5,631651212 | 0 | 1991,122556 | 46,71010234 | 134,3217459 |
| yp49_F04 | CroG8H-pENO2-pTPI1-AanCPR | 3,810548696 | 0 | 620,0036949 | 38,73805153 | 131,8116972 |
| yp50_D07 | SmusG8H-pENO2-pCCW12-RseCPR | 6,84239459 | 0 | 925,3659031 | 43,08937181 | 123,9098046 |
| yp51_A01 | OeuG8H-pENO2-pCCW12-CroCPR | 4,37552641 | 0 | 2007,341661 | 41,828541 | 122,2104943 |
| yp50_F10 | RsepG8H-pENO2-pCCW12-CloCPR | 4,212129207 | 0 | 983,4168456 | 34,93819477 | 100,4699002 |
| yp50_D05 | SmusG8H-pENO2-pURE2-CloCPR | 9,941620985 | 0 | 927,3319498 | 28,85710651 | 82,98283958 |
| yp50_A08 | VminG8H-pENO2-pCCW12-AraCPR | 2,633948973 | 0 | 1294,475665 | 28,68411897 | 82,48538855 |
| yp49_F05 | CroG8H-pENO2-pTPI1-AanCPR | 3,550161381 | 0 | 487,1068905 | 23,04201142 | 78,40370156 |
| yp50_G01 | RsepG8H-pCYC1-pCCW12-RseCPR | 2,708525027 | 0 | 1881,510579 | 26,48686695 | 76,1668683 |
| yp50_E07 | SmusG8H-pCYC1-pTPI1-OeuCPR | 6,63255802 | 0 | 1840,694046 | 26,30122887 | 75,63303879 |
| yp50_E08 | SmusG8H-pCYC1-pCCW12-OeuCPR | 4,43888775 | 0 | 1054,470029 | 25,19065345 | 72,43941639 |
| yp50_E06 | SmusG8H-pENO2-pURE2-CacCPR | 4,393836774 | 0 | 972,2468543 | 22,41555308 | 64,45920851 |
| yp50_C03 | VminG8H-pENO2-pTPI1-OeuCPR | 2,277114606 | 0 | 904,2719573 | 19,84016624 | 57,05330615 |
| yp50_G10 | RsepG8H-pENO2-pURE2-CacCPR | 2,917454788 | 0 | 916,207109 | 19,21148765 | 55,24544872 |
| yp49_G11 | CroG8H-pCYC1-pTPI1-CacCPR | 0 | 0 | 754,6718013 | 15,2495731 | 51,88882847 |
| yp49_C09 | OpumG8H-pENO2-pCCW12-CroCPR | 0 | 0 | 641,1446012 | 11,28045121 | 38,38333008 |
| yp50_B03 | VminG8H-pCYC1-pCCW12-RseCPR | 0 | 0 | 1716,722906 | 12,83526396 | 36,90968288 |
| yp51_A06 | OeuG8H-pCYC1-pCCW12-AanCPR | 0 | 0 | 2399,98713 | 12,43432535 | 36,32938206 |
| yp49_C11 | OpumG8H-pENO2-pCCW12-AanCPR | 1,289289923 | 0 | 617,7970603 | 10,27062405 | 34,9472504 |
| yp50_C08 | SmusG8H-pRPL15B-pTPI1-CroCPR | 0 | 0 | 1546,676152 | 11,92257384 | 34,28510866 |
| yp50_D04 | SmusG8H-pRPL15B-pTPI1-CloCPR | 0 | 0 | 1239,812058 | 11,10799518 | 31,94266833 |
| yp50_D08 | SmusG8H-pRPL15B-pTPI1-RseCPR | 0 | 0 | 949,5580327 | 10,67889628 | 30,70873154 |
| yp49_F03 | CroG8H-pCYC1-pTPI1-CroCPR | 1,118868795 | 0 | 500,6806578 | 8,01430537 | 27,26980709 |
| yp49_F01 | CroG8H-pPCK1-pTPI1-CroCPR | 1,350409857 | 0 | 621,345993 | 7,365141263 | 25,06093444 |
| yp50_C07 | SmusG8H-pRPL15B-pTPI1-CroCPR | 0 | 0 | 867,6266446 | 7,831343649 | 22,52017656 |
| yp49_F08 | CroG8H-pCYC1-pURE2-AraCPR | 1,253640043 | 0 | 882,8975409 | 5,739600807 | 19,52980322 |
| yp50_B02 | VminG8H-pENO2-pURE2-RseCPR | 0 | 0 | 1842,539358 | 6,230322106 | 17,91620444 |
| yp50_D02 | SmusG8H-pRPL15B-pCCW12-AraCPR | 0 | 0 | 1758,5958 | 6,165759466 | 17,73054511 |
| yp50_D06 | SmusG8H-pENO2-pURE2-CloCPR | 0 | 0 | 1438,102884 | 5,940724127 | 17,08342301 |
| yp51_C06 | OeuG8H-pENO2-pCCW12-CpoCPR | 0 | 0 | 2615,595356 | 4,794098799 | 14,00692374 |
| yp50_D01 | SmusG8H-pRPL15B-pCCW12-AraCPR | 0 | 0 | 1473,266739 | 4,683755207 | 13,46882463 |
| yp51_A08 | OeuG8H-pCYC1-pTPI1-AraCPR | 0 | 0 | 1591,876894 | 4,605194414 | 13,45500159 |
| yp50_F05 | RsepG8H-pCYC1-pURE2-AanCPR | 0 | 0 | 868,4137436 | 4,656776674 | 13,39124391 |
| yp50_C12 | SmusG8H-pRPL15B-pCCW12-AanCPR | 0 | 0 | 925,4692797 | 4,084635392 | 11,74596779 |
| yp50_A02 | VminG8H-pRPL15B-pTPI1-CroCPR | 0 | 0 | 1573,648146 | 3,229914997 | 9,288094008 |
| yp50_A07 | VminG8H-pRPL15B-pTPI1-AraCPR | 0 | 0 | 1693,097385 | 3,171907571 | 9,121285152 |
| yp50_F04 | RsepG8H-pCYC1-pURE2-AanCPR | 0 | 0 | 962,4212021 | 3,047825923 | 8,764470185 |
| yp51_B11 | OeuG8H-pPCK1-pURE2-CacCPR | 0 | 0 | 2237,832356 | 2,350935786 | 6,868731673 |
| yp49_E10 | OpumG8H-pENO2-pTPI1-CpoCPR | 0 | 0 | 765,539814 | 2,014252596 | 6,853779235 |
| yp51_E07 | CcalG8H-pCYC1-pURE2-OeuCPR | 0 | 0 | 2569,233123 | 2,235944362 | 6,532761103 |
| yp49_E04 | OpumG8H-pCYC1-pTPI1-CacCPR | 0 | 0 | 547,5282549 | 1,886535236 | 6,419202861 |
| yp50_B11 | VminG8H-pCYC1-pURE2-CacCPR | 0 | 0 | 1178,490707 | 2,036817008 | 5,857165859 |
| yp50_H01 | RsepG8H-pRPL15B-pCCW12-OeuCPR | 0 | 0 | 1135,510061 | 1,654537573 | 4,75786531 |
| yp49_F12 | CroG8H-pCYC1-pURE2-CloCPR | 0 | 0 | 790,9465298 | 1,325868406 | 4,511454704 |
| yp50_F07 | RsepG8H-pRPL15B-pCCW12-AraCPR | 0 | 0 | 1310,344177 | 1,547986128 | 4,451461012 |
| yp49_G01 | CroG8H-pENO2-pMLS1-RseCPR | 0 | 0 | 722,1177319 | 1,112403641 | 3,785110661 |
| yp50_G11 | RsepG8H-pRPL15B-pURE2-CacCPR | 0 | 0 | 2163,800169 | 1,24752813 | 3,587449998 |
| yp50_C10 | SmusG8H-pPCK1-pCCW12-AanCPR | 0 | 0 | 1299,060667 | 1,226744822 | 3,527684549 |
| yp51_C09 | CcalG8H-pENO2-pTPI1-CroCPR | 0 | 0 | 955,8828299 | 1,198360254 | 3,501250473 |
| yp51_A07 | OeuG8H-pCYC1-pURE2-AraCPR | 0 | 0 | 1605,325683 | 1,190719415 | 3,478926225 |
| yp51_A05 | OeuG8H-pRPL15B-pTPI1-AanCPR | 0 | 0 | 1274,656235 | 1,174344573 | 3,431083831 |
| yp50_B10 | VminG8H-pRPL15B-pURE2-CacCPR | 0 | 0 | 1994,055243 | 1,188093937 | 3,416538264 |
| yp50_A12 | VminG8H-pRPL15B-pCCW12-CloCPR | 0 | 0 | 987,0032206 | 1,115554969 | 3,207941829 |
| yp50_G04 | RsepG8H-pENO2-pTPI1-AhuCPR | 0 | 0 | 1394,199969 | 1,070810638 | 3,079272947 |
| yp50_C02 | VminG8H-pRPL15B-pCCW12-OeuCPR | 0 | 0 | 877,9359069 | 0,947957857 | 2,725991767 |
| yp51_D06 | CcalG8H-pRPL15B-pCCW12-CloCPR | 0 | 0 | 1112,254924 | 0,880562414 | 2,572740174 |
| yp49_G12 | CroG8H-pCYC1-pMLS1-CacCPR | 0 | 0 | 681,6958245 | 0,715858416 | 2,435809469 |
| yp49_F09 | CroG8H-pPCK1-pTPI1-AraCPR | 0 | 0 | 569,5059418 | 0,709600995 | 2,41451771 |
| yp50_H06 | RsepG8H-pCYC1-pTPI1-CpoCPR | 0 | 0 | 1544,592885 | 0,808004599 | 2,32353566 |
| yp51_B01 | OeuG8H-pRPL15B-pMLS1-RseCPR | 0 | 0 | 983,2773873 | 0,748061519 | 2,185612163 |
| yp51_A04 | OeuG8H-pPCK1-pCCW12-AanCPR | 0 | 0 | 1314,192011 | 0,686727482 | 2,00641244 |
| yp50_A05 | VminG8H-pRPL15B-pTPI1-AanCPR | 0 | 0 | 1076,92099 | 0,634958629 | 1,825916609 |
| yp49_F11 | CroG8H-pRPL15B-pMLS1-CloCPR | 0 | 0 | 937,7985145 | 0,474308762 | 1,613902621 |
| yp51_C01 | OeuG8H-pRPL15B-pTPI1-OeuCPR | 0 | 0 | 1078,351641 | 0,4431894 | 1,294866958 |
| yp49_D07 | OpumG8H-pENO2-pMLS1-RseCPR | 0 | 0 | 962,7378542 | 0,346186177 | 1,177947411 |
| yp50_G09 | RsepG8H-pENO2-pURE2-AniCPR | 0 | 0 | 894,9721508 | 0,374555667 | 1,077089722 |
| yp50_A03 | VminG8H-pRPL15B-pURE2-CroCPR | 0 | 0 | 1939,702053 | 0,374048665 | 1,075631763 |
| yp49_A06 | CacuG8H-pENO2-pTPI1-AanCPR | 0 | 0 | 1061,010835 | 0,315967733 | 1,075124883 |
| yp49_D02 | OpumG8H-pRPL15B-pCCW12-AraCPR | 0 | 0 | 761,0185433 | 0,291422432 | 0,9916060264 |
| yp50_F01 | RsepG8H-pPCK1-pCCW12-CroCPR | 0 | 0 | 1073,700811 | 0,324040183 | 0,9318250433 |
| yp49_A07 | CacuG8H-pCYC1-pCCW12-AraCPR | 0 | 0 | 756,5657284 | 0,273532239 | 0,9307321154 |
| yp49_A01 | CacuG8H-pENO2-pCCW12-CroCPR | 0 | 0 | 916,8168206 | 0,271198899 | 0,9227925962 |
| yp49_D05 | OpumG8H-pRPL15B-pCCW12-CloCPR | 0 | 0 | 498,8517726 | 0,271036809 | 0,922241062 |
| yp49_E11 | OpumG8H-pENO2-pMLS1-CpoCPR | 0 | 0 | 883,370915 | 0,270060501 | 0,918919036 |
| yp49_B12 | CacuG8H-pENO2-pTPI1-CacCPR | 0 | 0 | 1128,980271 | 0,263737113 | 0,8974028144 |
| yp49_F07 | CroG8H-pPCK1-pURE2-AraCPR | 0 | 0 | 669,1105418 | 0,256488305 | 0,8727377203 |
| yp49_C10 | OpumG8H-pRPL15B-pCCW12-AanCPR | 0 | 0 | 727,2523218 | 0,250470249 | 0,8522604339 |
| yp49_E09 | OpumG8H-pRPL15B-pURE2-OeuCPR | 0 | 0 | 1088,080352 | 0,227093503 | 0,7727177506 |
| yp49_G05 | CroG8H-pRPL15B-pURE2-AhuCPR | 0 | 0 | 772,0393709 | 0,209972294 | 0,7144604164 |
| yp51_B03 | OeuG8H-pRPL15B-pCCW12-RseCPR | 0 | 0 | 1016,539271 | 0,243293062 | 0,710829607 |
| yp51_E08 | CcalG8H-pRPL15B-pCCW12-OeuCPR | 0 | 0 | 994,3081201 | 0,219880506 | 0,6424251163 |
| yp50_C01 | VminG8H-pRPL15B-pMLS1-OeuCPR | 0 | 0 | 899,8726376 | 0,21651243 | 0,6226132284 |
| yp51_D12 | CcalG8H-pENO2-pMLS1-AhuCPR | 0 | 0 | 1711,361528 | 0,202342763 | 0,5911850733 |
| yp49_F02 | CroG8H-pRPL15B-pCCW12-CroCPR | 0 | 0 | 552,4335663 | 0,152172192 | 0,5177873975 |
| yp49_C01 | CacuG8H-pENO2-pCCW12-OeuCPR | 0 | 0 | 853,1795157 | 0,149625874 | 0,5091231905 |
| yp51_E12 | CcalG8H-pCYC1-pTPI1-CpoCPR | 0 | 0 | 1103,389383 | 0,15355561 | 0,4486435947 |
| yp49_H01 | CroG8H-pPCK1-pURE2-OeuCPR | 0 | 0 | 706,5267195 | 0,128271482 | 0,4364618526 |
| yp49_C03 | CacuG8H-pPCK1-pTPI1-OeuCPR | 0 | 0 | 950,9398202 | 0,125184236 | 0,4259570616 |
| yp50_G05 | RsepG8H-pCYC1-pMLS1-AhuCPR | 0 | 0 | 859,1142242 | 0,146244605 | 0,4205477979 |
| yp49_E01 | OpumG8H-pCYC1-pCCW12-AniCPR | 0 | 0 | 766,6255773 | 0,096245547 | 0,3274890809 |
| yp49_G09 | CroG8H-pRPL15B-pTPI1-AniCPR | 0 | 0 | 524,957627 | 0,091211339 | 0,3103594765 |
| yp50_A10 | VminG8H-pRPL15B-pURE2-CloCPR | 0 | 0 | 1389,879441 | 0,101051699 | 0,2905889724 |
| yp49_F10 | CroG8H-pPCK1-pTPI1-CloCPR | 0 | 0 | 463,2599925 | 0,077171721 | 0,2625876913 |
| yp49_B11 | CacuG8H-pCYC1-pURE2-CacCPR | 0 | 0 | 1223,612551 | 0,076985129 | 0,2619527856 |
| yp49_A02 | CacuG8H-pPCK1-pURE2-CroCPR | 0 | 0 | 1003,256653 | 0,075111672 | 0,2555780833 |
| yp49_B02 | CacuG8H-pENO2-pURE2-RseCPR | 0 | 0 | 1336,272488 | 0,071349681 | 0,2427773771 |
| yp49_A08 | CacuG8H-pCYC1-pURE2-AraCPR | 0 | 0 | 631,2707215 | 0,062366888 | 0,2122121539 |
| yp49_F06 | CroG8H-pPCK1-pMLS1-AanCPR | 0 | 0 | 885,1671701 | 0,060069359 | 0,2043944867 |
| yp49_A04 | CacuG8H-pPCK1-pURE2-AanCPR | 0 | 0 | 848,287124 | 0,056514692 | 0,1922992296 |
| yp49_D09 | OpumG8H-pPCK1-pURE2-RseCPR | 0 | 0 | 681,4642055 | 0,052854477 | 0,1798448306 |
| yp49_D01 | OpumG8H-pRPL15B-pMLS1-AraCPR | 0 | 0 | 746,4809637 | 0,049041597 | 0,1668709673 |
| yp49_C04 | CacuG8H-pCYC1-pCCW12-CpoCPR | 0 | 0 | 1195,336587 | 0,047430122 | 0,1613876958 |
| yp49_C02 | CacuG8H-pPCK1-pURE2-OeuCPR | 0 | 0 | 937,5099488 | 0,047390542 | 0,1612530193 |
| yp51_E04 | CcalG8H-pCYC1-pMLS1-CacCPR | 0 | 0 | 946,1147173 | 0,053133423 | 0,1552399804 |
| yp51_D05 | CcalG8H-pPCK1-pURE2-CloCPR | 0 | 0 | 2132,578642 | 0 | 0 |
| yp51_E05 | CcalG8H-pPCK1-pTPI1-CacCPR | 0 | 0 | 1098,071795 | 0 | 0 |
| yp49_B03 | CacuG8H-pCYC1-pURE2-RseCPR | 0 | 0 | 920,5952191 | 0 | 0 |
| yp51_D11 | CcalG8H-pRPL15B-pMLS1-AhuCPR | 0 | 0 | 1133,414992 | 0 | 0 |
| yp51_C05 | OeuG8H-pCYC1-pURE2-CpoCPR | 0 | 0 | 1175,46986 | 0 | 0 |
| yp51_B05 | OeuG8H-pCYC1-pMLS1-AhuCPR | 0 | 0 | 1089,844046 | 0 | 0 |
| yp51_C04 | OeuG8H-pRPL15B-pURE2-CpoCPR | 0 | 0 | 2771,931563 | 0 | 0 |
| yp49_E03 | OpumG8H-pRPL15B-pMLS1-AniCPR | 0 | 0 | 715,084384 | 0 | 0 |
| yp51_B06 | OeuG8H-pCYC1-pURE2-AhuCPR | 0 | 0 | 968,9137374 | 0 | 0 |
| yp51_D07 | CcalG8H-pRPL15B-pURE2-RseCPR | 0 | 0 | 2273,398103 | 0 | 0 |
| yp50_E02 | SmusG8H-pRPL15B-pCCW12-AniCPR | 0 | 0 | 1909,716721 | 0 | 0 |
| yp51_C07 | CcalG8H-pCYC1-pMLS1-CroCPR | 0 | 0 | 1062,858395 | 0 | 0 |
| yp51_C12 | CcalG8H-pPCK1-pMLS1-AanCPR | 0 | 0 | 1130,602453 | 0 | 0 |
| yp51_C11 | CcalG8H-pRPL15B-pMLS1-AanCPR | 0 | 0 | 1604,158931 | 0 | 0 |
| yp51_D08 | CcalG8H-pENO2-pMLS1-RseCPR | 0 | 0 | 1081,669636 | 0 | 0 |
| yp51_A03 | OeuG8H-pRPL15B-pMLS1-CroCPR | 0 | 0 | 1010,43009 | 0 | 0 |
| yp51_D09 | CcalG8H-pRPL15B-pMLS1-RseCPR | 0 | 0 | 1174,80925 | 0 | 0 |
| yp51_A10 | OeuG8H-pRPL15B-pURE2-CloCPR | 0 | 0 | 2106,860879 | 0 | 0 |
| yp51_A12 | OeuG8H-pRPL15B-pMLS1-CloCPR | 0 | 0 | 1804,613775 | 0 | 0 |
| yp51_C10 | CcalG8H-pPCK1-pURE2-AanCPR | 0 | 0 | 2157,667776 | 0 | 0 |
| yp51_D10 | CcalG8H-pRPL15B-pCCW12-AhuCPR | 0 | 0 | 2235,718855 | 0 | 0 |
| yp51_E11 | CcalG8H-pPCK1-pMLS1-CpoCPR | 0 | 0 | 1567,396099 | 0 | 0 |
| yp50_A04 | VminG8H-pPCK1-pMLS1-AanCPR | 0 | 0 | 1208,291917 | 0 | 0 |
| yp50_F12 | RsepG8H-pRPL15B-pMLS1-CloCPR | 0 | 0 | 1683,887779 | 0 | 0 |
| yp51_E02 | CcalG8H-pCYC1-pURE2-AniCPR | 0 | 0 | 1208,84877 | 0 | 0 |
| yp50_G06 | RsepG8H-pRPL15B-pTPI1-AhuCPR | 0 | 0 | 851,7897266 | 0 | 0 |
| yp50_E03 | SmusG8H-pRPL15B-pMLS1-AniCPR | 0 | 0 | 1938,499214 | 0 | 0 |
| yp49_E07 | OpumG8H-pPCK1-pMLS1-OeuCPR | 0 | 0 | 1067,303745 | 0 | 0 |
| yp50_B08 | VminG8H-pRPL15B-pCCW12-AniCPR | 0 | 0 | 1305,104406 | 0 | 0 |
| yp50_G07 | RsepG8H-pCYC1-pCCW12-AniCPR | 0 | 0 | 831,4886344 | 0 | 0 |
| yp49_A09 | CacuG8H-pPCK1-pMLS1-AraCPR | 0 | 0 | 691,2610411 | 0 | 0 |
| yp50_B07 | VminG8H-pPCK1-pURE2-AniCPR | 0 | 0 | 1180,834359 | 0 | 0 |
| yp50_F06 | RsepG8H-pPCK1-pURE2-AanCPR | 0 | 0 | 1417,717145 | 0 | 0 |
| yp50_F08 | RsepG8H-pRPL15B-pMLS1-AraCPR | 0 | 0 | 1380,345188 | 0 | 0 |
| yp49_B10 | CacuG8H-pCYC1-pURE2-CacCPR | 0 | 0 | 681,3554238 | 0 | 0 |
| yp50_B06 | VminG8H-pPCK1-pURE2-AhuCPR | 0 | 0 | 1728,399819 | 0 | 0 |
| yp49_G10 | CroG8H-pPCK1-pMLS1-CacCPR | 0 | 0 | 715,1072525 | 0 | 0 |
| yp50_B05 | VminG8H-pCYC1-pURE2-AhuCPR | 0 | 0 | 1482,311528 | 0 | 0 |
| yp50_E04 | SmusG8H-pPCK1-pMLS1-CacCPR | 0 | 0 | 1090,798003 | 0 | 0 |
| yp50_C04 | VminG8H-pRPL15B-pCCW12-CpoCPR | 0 | 0 | 846,1942953 | 0 | 0 |
| yp49_D12 | OpumG8H-pRPL15B-pTPI1-AhuCPR | 0 | 0 | 619,3532705 | 0 | 0 |
| yp49_E12 | OpumG8H-pPCK1-pURE2-CpoCPR | 0 | 0 | 568,424716 | 0 | 0 |
| yp51_D02 | CcalG8H-pRPL15B-pMLS1-AraCPR | 0 | 0 | 1004,379815 | 0 | 0 |
| yp50_E12 | SmusG8H-pRPL15B-pCCW12-CpoCPR | 0 | 0 | 1742,491768 | 0 | 0 |
| yp51_C02 | OeuG8H-pPCK1-pCCW12-OeuCPR | 0 | 0 | 1018,289695 | 0 | 0 |
| yp51_B02 | OeuG8H-pPCK1-pMLS1-RseCPR | 0 | 0 | 1100,537442 | 0 | 0 |
| yp50_G02 | RsepG8H-pPCK1-pURE2-RseCPR | 0 | 0 | 1160,021864 | 0 | 0 |
| yp49_B04 | CacuG8H-pCYC1-pCCW12-AhuCPR | 0 | 0 | 841,0308888 | 0 | 0 |
| yp50_G12 | RsepG8H-pPCK1-pMLS1-CacCPR | 0 | 0 | 1645,151979 | 0 | 0 |
| yp50_B04 | VminG8H-pPCK1-pURE2-AhuCPR | 0 | 0 | 1108,870809 | 0 | 0 |
| yp50_D12 | SmusG8H-pRPL15B-pURE2-AhuCPR | 0 | 0 | 1579,821396 | 0 | 0 |
| yp49_B06 | CacuG8H-pPCK1-pCCW12-AhuCPR | 0 | 0 | 731,4989764 | 0 | 0 |
| yp49_D04 | OpumG8H-pPCK1-pURE2-CloCPR | 0 | 0 | 872,7746411 | 0 | 0 |
| yp50_B12 | VminG8H-pCYC1-pMLS1-CacCPR | 0 | 0 | 1233,277094 | 0 | 0 |
| yp50_E01 | SmusG8H-pPCK1-pCCW12-AniCPR | 0 | 0 | 1322,633529 | 0 | 0 |
| yp50_D11 | SmusG8H-pRPL15B-pURE2-AhuCPR | 0 | 0 | 1129,826177 | 0 | 0 |
| yp49_E05 | OpumG8H-pPCK1-pMLS1-CacCPR | 0 | 0 | 728,3260392 | 0 | 0 |
| yp49_H05 | CroG8H-pPCK1-pMLS1-CpoCPR | 0 | 0 | 676,327275 | 0 | 0 |
| yp49_B05 | CacuG8H-pPCK1-pURE2-AhuCPR | 0 | 0 | 672,2711334 | 0 | 0 |

**Table 3. Sorted strain performance of all fully genotyped strains in the second DBTL round.**

| **plate-well** | **names** | **Amt_uM_Loganin** | **Amt_uM_Secologanin** | **Amt_uM_Tryptamine** | **Amt_uM_Strictosidine** | **normalized_**  **strictosidine(%)** |
| --- | --- | --- | --- | --- | --- | --- |
| yp53-D09 | SmusG8H-pENO2-pTPI1-RseCPR | 7,054979763 | 0 | 774,9648421 | 47,69202744 | 169,6313959 |
| yp53-C06 | RsepG8H-pENO2-pTPI1-CloCPR | 8,910505149 | 0 | 942,4938148 | 44,84855721 | 159,5177175 |
| yp53-A07 | SmusG8H-pENO2-pTPI1-AanCPR | 7,860359137 | 0 | 820,2152727 | 44,71979192 | 159,0597241 |
| yp53-E02 | RsepG8H-pENO2-pTPI1-RseCPR | 5,506114555 | 0 | 924,4804019 | 40,80980643 | 145,1526555 |
| yp53-B02 | CroG8H-pENO2-pTPI1-AanCPR | 5,139113172 | 0 | 830,1376607 | 40,76194282 | 144,9824138 |
| yp53-C01 | SmusG8H-pCYC1-pCCW12-CloCPR | 5,944782007 | 0 | 1123,437012 | 40,34055827 | 143,4836298 |
| yp54-B01 | CroG8H-pENO2-pTPI1-CroCPR | 3,916937343 | 0 | 968,8466421 | 39,04466894 | 138,2677567 |
| yp53-E05 | CroG8H-pENO2-pCCW12-RseCPR | 4,067949178 | 0 | 792,0549682 | 37,29660365 | 132,6568669 |
| yp53-A10 | RsepG8H-pENO2-pCCW12-AanCPR | 3,391072801 | 0 | 846,3520846 | 36,46977402 | 129,7159925 |
| yp53-B05 | OeuG8H-pENO2-pTPI1-AanCPR | 4,699779702 | 0 | 994,698889 | 35,69348625 | 126,9548858 |
| yp53-C07 | RsepG8H-pENO2-pTPI1-CloCPR | 4,956388647 | 0 | 832,1421358 | 35,60440093 | 126,6380264 |
| yp53-C02 | SmusG8H-pCYC1-pTPI1-CloCPR | 4,185685742 | 0 | 979,9955748 | 35,0689843 | 124,7336521 |
| yp53-A06 | SmusG8H-pCYC1-pCCW12-AanCPR | 2,838787848 | 0 | 783,4952066 | 34,23506795 | 121,7675716 |
| yp54-C10 | CroG8H-pENO2-pCCW12-AraCPR | 3,58547569 | 0 | 883,5363126 | 32,85060784 | 116,3329073 |
| yp54-B05 | OeuG8H-pENO2-pTPI1-CroCPR | 2,940680875 | 0 | 1026,468 | 32,74180598 | 115,9476104 |
| yp54-A12 | RsepG8H-pENO2-pCCW12-CroCPR | 2,28627235 | 0 | 828,0913556 | 32,42157303 | 114,8135787 |
| yp54-A11 | RsepG8H-pENO2-pCCW12-CroCPR | 3,157059788 | 0 | 899,7981276 | 32,41057353 | 114,7746265 |
| yp54-A09 | RsepG8H-pENO2-pTPI1-CroCPR | 4,384179517 | 0 | 901,6366184 | 32,39656973 | 114,7250352 |
| yp54-A05 | SmusG8H-pENO2-pTPI1-CroCPR | 5,725198525 | 0 | 866,9232181 | 31,95842961 | 113,1734623 |
| yp53-D10 | SmusG8H-pCYC1-pTPI1-RseCPR | 5,280057892 | 0 | 1058,74296 | 30,9489304 | 110,0794105 |
| yp53-D06 | VminG8H-pENO2-pTPI1-RseCPR | 4,023233772 | 0 | 932,8833042 | 30,73676235 | 109,3247694 |
| yp53-E04 | RsepG8H-pENO2-pTPI1-RseCPR | 2,847971208 | 0 | 660,8714767 | 29,81386433 | 106,0421981 |
| yp54-D04 | OeuG8H-pENO2-pTPI1-AraCPR | 1,999756647 | 0 | 944,2784733 | 28,63213148 | 101,394139 |
| yp53-D08 | VminG8H-pENO2-pCCW12-RseCPR | 2,293022184 | 0 | 933,4301086 | 27,74011158 | 98,66625726 |
| yp53-E12 | OeuG8H-pENO2-pTPI1-RseCPR | 3,721896614 | 0 | 919,9040756 | 27,6921933 | 98,49582113 |
| yp54-C04 | SmusG8H-pENO2-pTPI1-AraCPR | 3,457464522 | 0 | 871,5535701 | 26,63404075 | 94,31835805 |
| yp54-C07 | RsepG8H-pENO2-pTPI1-AraCPR | 5,604359855 | 0 | 866,4201972 | 26,28236802 | 93,07298959 |
| yp54-D02 | OeuG8H-pENO2-pTPI1-AraCPR | 1,77796506 | 0 | 966,3462739 | 26,06058955 | 92,28761191 |
| yp53-A08 | SmusG8H-pCYC1-pTPI1-AanCPR | 2,683838824 | 0 | 749,8241168 | 24,75364985 | 88,04398559 |
| yp54-A03 | VminG8H-pENO2-pTPI1-CroCPR | 2,483075465 | 0 | 947,1655162 | 24,80108374 | 87,82736042 |
| yp54-A07 | SmusG8H-pENO2-pCCW12-CroCPR | 7,57724781 | 0 | 974,620347 | 24,76383324 | 87,6954463 |
| yp54-C01 | SmusG8H-pCYC1-pTPI1-AraCPR | 2,396037195 | 0 | 842,0344901 | 23,6663117 | 83,80882502 |
| yp53-A04 | VminG8H-pENO2-pCCW12-AanCPR | 1,50126031 | 0 | 823,8000874 | 22,59774959 | 80,3758618 |
| yp53-B08 | OeuG8H-pENO2-pCCW12-AanCPR | 1,5479584 | 0 | 740,1410436 | 22,49698249 | 80,01745255 |
| yp53-B06 | OeuG8H-pENO2-pCCW12-AanCPR | 0 | 0 | 783,9396143 | 21,59328835 | 76,80318579 |
| yp53-C03 | SmusG8H-pCYC1-pTPI1-CloCPR | 1,797113069 | 0 | 767,6454625 | 21,19806246 | 75,39744309 |
| yp53-D11 | SmusG8H-pCYC1-pTPI1-RseCPR | 3,504055023 | 0 | 929,7938959 | 21,07241646 | 74,95054436 |
| yp54-C12 | CroG8H-pCYC1-pCCW12-AraCPR | 0 | 0 | 972,5675072 | 19,31132744 | 68,38664524 |
| yp54-B10 | VminG8H-pENO2-pCCW12-AraCPR | 0,763484297 | 0 | 814,5903383 | 18,6488437 | 66,04061073 |
| yp53-B04 | CroG8H-pCYC1-pTPI1-AanCPR | 1,938178185 | 0 | 1046,657914 | 18,07923742 | 64,30438051 |
| yp54-B08 | OeuG8H-pENO2-pTPI1-CroCPR | 2,928218659 | 0 | 982,904756 | 16,01610344 | 56,71736381 |
| yp53-E03 | RsepG8H-pCYC1-pCCW12-RseCPR | 0 | 0 | 940,88909 | 15,75954539 | 56,0536808 |
| yp53-A12 | RsepG8H-pCYC1-pTPI1-AanCPR | 2,128574722 | 0 | 1070,432162 | 15,44535688 | 54,93617252 |
| yp54-F05 | VminG8H-pENO2-pCCW12-AanCPR | 0,994538655 | 0 | 890,2598802 | 15,39707673 | 54,52522243 |
| yp53-A09 | RsepG8H-pCYC1-pTPI1-AanCPR | 1,294812747 | 0 | 907,5359256 | 15,31187214 | 54,4613929 |
| yp54-C02 | SmusG8H-pENO2-pTPI1-AraCPR | 4,255545878 | 0 | 937,1612096 | 15,04377179 | 53,27407386 |
| yp53-C08 | RsepG8H-pCYC1-pTPI1-CloCPR | 1,583715867 | 0 | 1123,649906 | 14,74325725 | 52,43893877 |
| yp53-C09 | CroG8H-pCYC1-pTPI1-CloCPR | 1,913996619 | 0 | 1050,590401 | 14,61778276 | 51,99265007 |
| yp53-E07 | CroG8H-pCYC1-pCCW12-RseCPR | 0 | 0 | 791,0259144 | 14,18377971 | 50,44898445 |
| yp54-B04 | CroG8H-pCYC1-pTPI1-CroCPR | 0 | 0 | 970,3137686 | 14,05958497 | 49,78880155 |
| yp53-C05 | RsepG8H-pCYC1-pCCW12-CloCPR | 0 | 0 | 1094,24756 | 13,94506097 | 49,59990767 |
| yp54-C05 | RsepG8H-pCYC1-pTPI1-AraCPR | 1,120564578 | 0 | 1084,159381 | 13,89445125 | 49,20401828 |
| yp53-B09 | VminG8H-pENO2-pTPI1-CloCPR | 2,096557201 | 0 | 983,312202 | 13,81069039 | 49,12197729 |
| yp53-C10 | CroG8H-pCYC1-pTPI1-CloCPR | 0 | 0 | 975,1446261 | 13,47359444 | 47,92299164 |
| yp54-C09 | CroG8H-pCYC1-pTPI1-AraCPR | 2,023879297 | 0 | 859,3435027 | 13,12227039 | 46,46951654 |
| yp53-A11 | RsepG8H-pCYC1-pTPI1-AanCPR | 0 | 0 | 773,5169829 | 11,88336506 | 42,26685069 |
| yp53-E01 | RsepG8H-pCYC1-pCCW12-RseCPR | 0 | 0 | 899,5546186 | 11,75307123 | 41,80342052 |
| yp54-A10 | RsepG8H-pCYC1-pTPI1-CroCPR | 0 | 0 | 926,9069467 | 10,79119438 | 38,2145445 |
| yp53-D05 | VminG8H-pENO2-pCCW12-RseCPR | 0 | 0 | 875,8538073 | 9,272851839 | 32,98175577 |
| yp53-A02 | VminG8H-pCYC1-pCCW12-AanCPR | 0 | 0 | 913,3709712 | 7,921695535 | 28,1759519 |
| yp53-B10 | VminG8H-pCYC1-pCCW12-CloCPR | 0 | 0 | 821,1026276 | 7,335207418 | 26,08992614 |
| yp54-A02 | VminG8H-pCYC1-pCCW12-CroCPR | 0 | 0 | 994,3880851 | 6,808027216 | 24,10906984 |
| yp54-F06 | VminG8H-pCYC1-pCCW12-AanCPR | 0 | 0 | 834,7434403 | 5,878048137 | 20,81576183 |
| yp53-B07 | OeuG8H-pCYC1-pCCW12-AanCPR | 0 | 0 | 864,854188 | 5,583939914 | 19,86100346 |
| yp53-D04 | OeuG8H-pCYC1-pCCW12-CloCPR | 0 | 0 | 889,6035514 | 5,326136042 | 18,94404453 |
| yp53-B12 | VminG8H-pCYC1-pTPI1-CloCPR | 0 | 0 | 968,186712 | 5,241370601 | 18,64255011 |
| yp54-C11 | CroG8H-pCYC1-pTPI1-AraCPR | 0,686359445 | 0 | 887,3231931 | 4,812066268 | 17,04083107 |
| yp54-A01 | VminG8H-pCYC1-pCCW12-CroCPR | 0 | 0 | 1058,714386 | 4,711870253 | 16,68600982 |
| yp54-B09 | VminG8H-pCYC1-pTPI1-AraCPR | 0 | 0 | 848,7677131 | 4,708556835 | 16,67427611 |
| yp54-A06 | SmusG8H-pCYC1-pCCW12-CroCPR | 0,768868795 | 0 | 905,3955124 | 4,679736887 | 16,57221686 |
| yp54-B07 | OeuG8H-pENO2-pTPI1-CroCPR | 1,179642912 | 0 | 793,6625702 | 4,604644062 | 16,30629282 |
| yp54-A04 | VminG8H-pCYC1-pTPI1-CroCPR | 0 | 0 | 876,739115 | 4,326963454 | 15,32295051 |
| yp53-B11 | VminG8H-pCYC1-pTPI1-CloCPR | 0 | 0 | 826,926938 | 4,260934467 | 15,15532679 |
| yp53-E09 | OeuG8H-pCYC1-pTPI1-RseCPR | 0 | 0 | 1004,352709 | 4,184889835 | 14,88485062 |
| yp53-B01 | CroG8H-pCYC1-pTPI1-AanCPR | 0 | 0 | 1087,656035 | 3,905533671 | 13,89123432 |
| yp53-E10 | OeuG8H-pCYC1-pTPI1-RseCPR | 0 | 0 | 961,5276058 | 3,492197071 | 12,42107529 |
| yp53-D01 | OeuG8H-pCYC1-pTPI1-CloCPR | 0 | 0 | 854,1031457 | 3,163020902 | 11,25025879 |
| yp53-D03 | OeuG8H-pCYC1-pTPI1-CloCPR | 0 | 0 | 806,7269817 | 2,726746706 | 9,698515134 |
| yp53-A01 | VminG8H-pCYC1-pCCW12-AanCPR | 0 | 0 | 1123,859574 | 2,546823228 | 9,058561826 |
| yp54-D01 | OeuG8H-pCYC1-pCCW12-AraCPR | 0 | 0 | 862,2470291 | 2,488403038 | 8,812109695 |
| yp54-B06 | OeuG8H-pENO2-pCCW12-CroCPR | 0 | 0 | 930,1164024 | 2,327762595 | 8,243238341 |
| yp54-B11 | VminG8H-pCYC1-pTPI1-AraCPR | 0 | 0 | 796,0811197 | 1,799734248 | 6,373346831 |
| yp54-E01 | RsepG8H-pENO2-pTPI1-AhuCPR | 0 | 0 | 921,8252653 | 1,784198127 | 6,318329215 |
| yp54-E06 | CroG8H-pENO2-pCCW12-AhuCPR | 0 | 0 | 959,7219823 | 1,709109071 | 6,052418513 |
| yp53-F01 | VminG8H-pENO2-pCCW12-AniCPR | 0 | 0 | 1181,431781 | 1,694404131 | 6,026670564 |
| yp53-E06 | CroG8H-pCYC1-pTPI1-RseCPR | 0 | 0 | 1005,347335 | 1,611958837 | 5,733428463 |
| yp54-E05 | CroG8H-pENO2-pCCW12-AhuCPR | 0 | 0 | 819,6875796 | 1,501958271 | 5,318841377 |
| yp54-D06 | VminG8H-pENO2-pTPI1-AhuCPR | 0 | 0 | 1183,558158 | 1,448709878 | 5,130274383 |
| yp54-E11 | OeuG8H-pENO2-pTPI1-AhuCPR | 0 | 0 | 910,2848833 | 1,307859472 | 4,631484915 |
| yp53-F10 | RsepG8H-pENO2-pTPI1-AniCPR | 0 | 0 | 1108,332 | 1,297344365 | 4,614405119 |
| yp53-F07 | SmusG8H-pENO2-pCCW12-AniCPR | 0 | 0 | 1069,685239 | 1,116576512 | 3,971448531 |
| yp54-D12 | SmusG8H-pCYC1-pTPI1-AhuCPR | 0 | 0 | 922,1936025 | 1,049919897 | 3,71805096 |
| yp54-E09 | OeuG8H-pENO2-pTPI1-AhuCPR | 0 | 0 | 789,4004619 | 0,997638389 | 3,532907968 |
| yp54-E08 | CroG8H-pCYC1-pTPI1-AhuCPR | 0 | 0 | 1021,519529 | 0,919757996 | 3,257112385 |
| yp54-E10 | OeuG8H-pENO2-pCCW12-AhuCPR | 0 | 0 | 921,471845 | 0,848364966 | 3,004290313 |
| yp54-E04 | RsepG8H-pCYC1-pCCW12-AhuCPR | 0 | 0 | 1009,589352 | 0,843157359 | 2,985848765 |
| yp54-C06 | RsepG8H-pCYC1-pTPI1-AraCPR | 0 | 0 | 792,6186431 | 0,842966998 | 2,985174645 |
| yp54-D08 | VminG8H-pENO2-pCCW12-AhuCPR | 0 | 0 | 1129,718025 | 0,813730516 | 2,881640337 |
| yp54-F02 | OeuG8H-pENO2-pCCW12-AniCPR | 0 | 0 | 1007,713513 | 0,794647266 | 2,814061376 |
| yp54-B12 | VminG8H-pCYC1-pTPI1-AraCPR | 0 | 0 | 891,6029522 | 0,759764027 | 2,690530371 |
| yp54-E07 | CroG8H-pCYC1-pCCW12-AhuCPR | 0 | 0 | 924,1415117 | 0,745224947 | 2,639043547 |
| yp54-F03 | OeuG8H-pENO2-pCCW12-AniCPR | 0 | 0 | 1003,72886 | 0,723003562 | 2,560351599 |
| yp54-F04 | OeuG8H-pENO2-pTPI1-AniCPR | 0 | 0 | 838,9994008 | 0,637065797 | 2,256022678 |
| yp53-E08 | CroG8H-pCYC1-pCCW12-RseCPR | 0 | 0 | 774,2920859 | 0,529311872 | 1,882660824 |
| yp53-F09 | RsepG8H-pCYC1-pCCW12-AniCPR | 0 | 0 | 1054,366652 | 0,516182596 | 1,835962507 |
| yp53-E11 | OeuG8H-pCYC1-pCCW12-RseCPR | 0 | 0 | 776,3233741 | 0,477411086 | 1,698059681 |
| yp54-D05 | VminG8H-pENO2-pCCW12-AhuCPR | 0 | 0 | 975,650181 | 0,422228924 | 1,495227075 |
| yp53-B03 | CroG8H-pCYC1-pCCW12-AanCPR | 0 | 0 | 961,0746162 | 0,414028309 | 1,472619298 |
| yp53-D02 | OeuG8H-pCYC1-pCCW12-CloCPR | 0 | 0 | 1006,695581 | 0,326690163 | 1,161974262 |
| yp53-F08 | SmusG8H-pCYC1-pCCW12-AniCPR | 0 | 0 | 952,8527213 | 0,272293948 | 0,9684973561 |
| yp53-G02 | CroG8H-pCYC1-pTPI1-AniCPR | 0 | 0 | 1456,925634 | 0,24371525 | 0,8668484078 |
| yp53-F05 | SmusG8H-pENO2-pCCW12-AniCPR | 0 | 0 | 708,073811 | 0,202178789 | 0,7191111813 |
| yp53-F11 | RsepG8H-pCYC1-pTPI1-AniCPR | 0 | 0 | 925,7417551 | 0,093646456 | 0,3330824857 |
| yp54-B02 | CroG8H-pENO2-pTPI1-CroCPR | 0 | 0 | 770,6129197 | 0,066441751 | 0,2352882508 |
| yp53-F06 | SmusG8H-pCYC1-pTPI1-AniCPR | 0 | 0 | 818,5590501 | 0,012795673 | 0,04551175508 |
| yp54-D11 | SmusG8H-pENO2-pTPI1-AhuCPR | 0 | 0 | 1123,565741 | 0,000369414 | 0,001308195112 |
| yp54-D10 | SmusG8H-pCYC1-pTPI1-AhuCPR | 0 | 0 | 950,9413806 | 0 | 0 |
| yp53-F03 | VminG8H-pCYC1-pTPI1-AniCPR | 0 | 0 | 908,3735426 | 0 | 0 |
| yp53-D12 | SmusG8H-pENO2-pTPI1-RseCPR | 0 | 0 | 823,0233242 | 0 | 0 |
| yp53-F02 | VminG8H-pENO2-pCCW12-AniCPR | 0 | 0 | 552,0471851 | 0 | 0 |
| yp54-F01 | OeuG8H-pENO2-pCCW12-AniCPR | 0 | 0 | 1045,869448 | 0 | 0 |
| yp53-F04 | VminG8H-pCYC1-pCCW12-AniCPR | 0 | 0 | 897,7012108 | 0 | 0 |
| yp53-G04 | CroG8H-pCYC1-pCCW12-AniCPR | 0 | 0 | 1164,572251 | 0 | 0 |
| yp54-E12 | OeuG8H-pCYC1-pTPI1-AhuCPR | 0 | 0 | 929,876888 | 0 | 0 |
| yp53-G03 | CroG8H-pCYC1-pCCW12-AniCPR | 0 | 0 | 1320,74472 | 0 | 0 |
| yp54-D09 | SmusG8H-pCYC1-pTPI1-AhuCPR | 0 | 0 | 1147,896742 | 0 | 0 |
| yp53-F12 | RsepG8H-pCYC1-pTPI1-AniCPR | 0 | 0 | 919,9275808 | 0 | 0 |
| yp54-E02 | RsepG8H-pCYC1-pTPI1-AhuCPR | 1,298691019 | 0 | 995,709125 | 0 | 0 |
| yp53-G01 | CroG8H-pCYC1-pTPI1-AniCPR | 0 | 0 | 1236,868056 | 0 | 0 |
